# GAS2 encodes a 2-oxoglutarate dependent dioxygenase involved in ABA catabolism

**DOI:** 10.1101/2022.11.16.516706

**Authors:** Theo Lange, Nadiem Atiq, Maria João Pimenta Lange

## Abstract

Liu et al.^1^ recently reported the characterization of *Arabidopsis thaliana* GAS2 (Gain of Function in ABA-modulated Seed Germination 2), which was described as an enzyme that catalyzes the stereospecific hydration of GA_12_ to produce GA_12_ 16, 17-dihydro-16α-ol (DHGA_12_). However, as previously reported^2^, we did not find conversion of [17-^14^C]-labeled or [1-,7-,12-,18-^14^C_4_]-labeled GA_12_ by GAS2. Furthermore, the authors^1^ state the isolation of endogenous DHGA_12_ from dry Arabidopsis seeds, which we cannot confirm by our attempts to isolate this compound from 0.5 g dry Arabidopsis Col-0 seeds (data not shown). Instead, we present here data showing that the recombinant GAS2 enzyme is able to catabolize abscisic acid (ABA) to phaseic acid (PA) and further to a second product, putative 8’-carboxy-ABA (Fig. 1a).

---

ABA is known to be oxidative catabolized by cytochrome P450 monooxygenases^3^. Three different hydroxylation pathways have been described that oxidize one of the methyl groups of the ring structure, at C-7’, C-8’, or C-9’. 8’-hydroxylation appears to be the major pathway, and Arabidopsis *CYP707As* encode ABA 8’-hydroxylases that initiate the oxidation at C-8’ to produce 8’-hydroxy-ABA, which spontaneously autoisomerizes to PA^4,5^. This final cyclization step is reversible, with the final equilibrium being at PA^6^ (Fig. 1a).

**Fig. 1.**
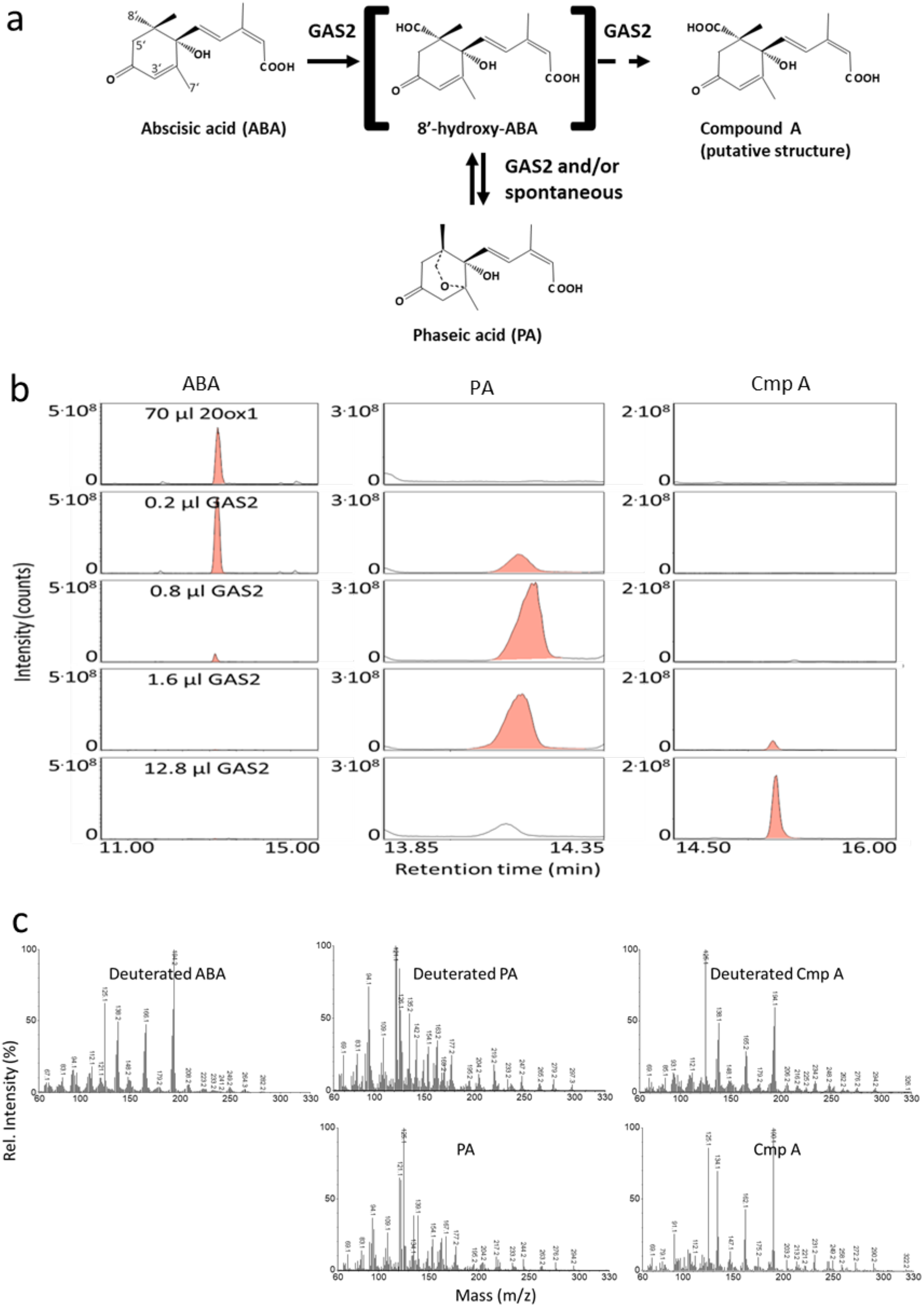
GAS2 is an ABA catabolizing oxidase. **(a)** The proposed ABA catabolic pathway catalyzed by GAS2. **(b)** Metabolism of d_6_-ABA incubated with 70 μl cell-lysate of recombinant ATGA2ox1 and cofactors (negative control, upper lane). Metabolism of d_6_-ABA incubated with different volumes of cell-lysates containing recombinant GAS2 and cofactors as described in Methods. Chromatograms of characteristic single ions are shown in the first row for ABA (194 Da), in the middle row for PA (125 DA), and in the right row for compound A (194 Da). Products that were identified on the basis of their mass spectra of the methyl ester derivatives are labelled in red. **(c)** Top lane: Representative mass spectra of the d_6_-ABA substrate (KRI = 2087), and of its incubation products, PA (KRI = 2141) and compound A (KRI = 2197), by recombinant GAS2 enzyme. Bottom lane: Mass spectra of unlabeled PA substrate (KRI = 2146), and of the incubation product, compound A (KRI = 2201), by recombinant GAS2 enzyme.

For this study, we expressed Arabidopsis GAS2 heterologously in *E. coli* according to our standard protocols^7,8^. As a control, we expressed Arabidopsis gibberellin 20-oxidase 1 (AtGA20ox1) in the same manner in *E. coli.* Both enzyme preparations were incubated with [^14^C]-labeled GA_12_ and deuterated ABA. As expected, recombinant AtGA20ox1 (70 μl cell-lysate) converts [^14^C]GA_12_ to a single product [^14^C]GA_9_. Surprisingly, in incubations with recombinant GAS2 [^14^C]GA_12_ is not converted^2^ (data not shown). In contrast, recombinant GAS2 completely converts deuterated ABA at 1.6 μl of cell lysate or more (Fig. 1b, left column). At low volumes, an intermediate product is formed first, which disappears at high lysate volumes (Fig. 1b, middle column) when a second product appears (compound A, Fig. 1b, right column). On the other hand, no products are formed in control incubations with AtGA20ox1 and deuterated ABA (Fig. 1b, upper lane), suggesting that the *E. coli* cell lysate is devoid of ABA-oxidizing activities.

Both the mass spectra and the KRI of the methyl esters of the intermediate product correspond to those of deuterated PA. The relatively low intensity of the *d4*-fragment ions in the mass spectrum of deuterated PA could indicate an isotope effect of GAS2 enzyme activity. However, there is no known ABA catabolite consistent with the mass spectrum of compound A. In fact, compound A shares several fragment ions with ABA^9^, suggesting structural similarities between the two compounds (Fig. 1c, spectra in the top lane). In addition, GAS2 also catalyzes the production of compound A from PA (Fig. 1c, spectra in the lower lane). Unlabeled compound A shows a molecular ion (M^+^) of 322 mass units in good agreement with ABA, containing an additional carboxy group (as a methyl ester) most likely at the 8’-position. This structure suggests that GAS2 catalyzes the conversion of PA to 8’-carboxy-ABA *via* 8’-hydroxy-ABA, and 8’-aldehyde-ABA. The structure for the latter is not indicated in Fig. 1a. Moreover, the intermediates have not been isolated, possibly because of their instability or rapid conversion^10^. Although such a stepwise oxidation pathway from methyl to the carboxyl group *via* the alcohol and aldehyde has been found in reactions catalyzed by other 2-oxoglutarate dependent dioxygenases, including gibberellin 20-oxidases^2^.

Xiong et al.^11^ observed that Arabidopsis *GAS2* overexpressing lines (here referred to as GIM2) show decreased endogenous ABA levels, while the Arabidopsis knock-out mutant shows increased endogenous ABA levels. Moreover, *GAS2* overexpressing lines show lower sensitivity to ABA in germination and early seedling development than WT and increased sensitivity to ABA in *GAS2* loss-of-function mutants^1^. Consistent with these observations and our own results, we conclude that *GAS2* encodes a 2-oxoglutarate-dependent dioxygenase that initiates a novel ABA catabolic pathway, rather than gibberellin anabolism as proposed by Liu et al.^1^

## Methods

### Heterologous expression of recombinant GAS2 and AtGA20ox1

*AtGAS2* and *AtGA20ox1* coding sequences were custom synthesized in pET21a expression vector (BioCat). Respective plasmid DNAs were used to transform BL21 Star^™^ *E. coli* (Invitrogen), according to the manufacturer’s instructions. Recombinant GAS2 and AtGA20ox1 were produced as previously described ^7,8^.

### Enzyme assays and analysis of incubation products

3’,5’,5’,7’,7’,7’-d_6_-labelled ABA was purchased from OlChemIm, Czech Republic. PA was a gift from Professor Eiji Nambara (University of Toronto, Canada). 17-^14^C-Labeled GA_12_ was a gift from Professor L. Mander (Canberra, Australia). Preparations of *E. coli* cell lysates were incubated with 2-oxoglutarate and ascorbate (100mM each, final concentrations). FeSO_4_ (0.5 mM), catalase (1mg/ml), and the substrates (2 μl in methanol for 17-^14^C-labeled GA_12_ (1 nmol), and 5 μl in methanol for 3’,5’,5’,7’,7’,7’-d_6_-labelled ABA (500 ng) and PA (500 ng)) were added in a total volume of 100 μl and incubated at 30°C for 16 h.

Incubation products were extracted and analyzed by reverse-phase HPLC with on-line radiocounting, as described previously^12^, using gradients of increasing methanol in acidic water, at 1 ml·min^-1^ as follows: 50% methanol, followed by five 0.5-min steps to 57.4%, 60.6%, 61.8%, 62.3%, 62.5%, one 5-min step to 62.7%, one 1-min step to 63.2% and seven 2-min steps to 64.3%, 67.2%, 70.5%, 74.9%, 81%, 89% and 100% methanol. The retention times for the [^14^C]-labeled GAs were 14.73 min for GA_9_ and GA_15_, 14:50 min for GA_24_, and 24.37 min for GA_12_. The deuterated-labeled PA and putative 8’-carboxylated ABA eluted between 3 and 6 min, and ABA between 6 and 9 min.

The dried HPLC-fractions were redissolved in 100 μl methanol and methylated with 100 μl ethereal diazomethane. The derivatized samples were analyzed using a Thermo Scientific MS system ISQ 7000 equipped with a Thermo Scientific Trace 1300 gas chromatograph. Samples (1-2 μl) were injected into an Agilent DB-5MS UI capillary column (30 m long, 0.25 mm i.d., 0.25 μm film thickness; Agilent, USA) at an oven temperature of 60°C. The split value (30:1) was open after 1 min, after which the temperature was increased by 45°C min^-1^ to 175°C and then with 4°C min^-1^ to 280°C. The He inlet was pneumatic pressure controlled at a constant flow rate of 1.2 ml·min^-1^, and the injector, transfer line, and source temperatures were 280, 280 and 250°C, respectively. Mass spectra were acquired from 5 to 30 min after injection at an electron energy of 45 eV from 60 to 660 atomic mass units at 0.2 s per scan. For determination of Kovats retention index (KRI), 0.5 μl solution of 1 cm^2^ parafilm in 2 ml hexane were co-injected.

## Author contributions

T. L., and M. J. P. L. designed the experiments; T. L. performed the GC-MS analysis and the corresponding evaluation; N. A. carried out the expression and enzyme assay of recombinant GAS2; T. L., and M. J. P. L. supervised the study; T. L., and M. J. P. L. wrote and edited the manuscript.

## Acknowledgements

We thank Professor Eiji Nambara (University of Toronto, Canada) for the generous gift of authentic PA.

## Additional information

Correspondence and requests for materials should be addressed to Theo Lange.

